# Self-grooming promotes social attraction in mice via chemosensory communication

**DOI:** 10.1101/2021.10.08.463701

**Authors:** Yun-Feng Zhang, Emma Janke, Janardhan P. Bhattarai, Daniel W. Wesson, Minghong Ma

## Abstract

Self-grooming is a stereotyped behavior displayed by nearly all animals. Among other established functions, self-grooming is implicated in social communication in some animals. However, whether self-grooming specifically influences behaviors of nearby individuals has not been directly tested, partly due to the technical challenge of inducing self-grooming in a reliable and temporally controllable manner. We recently found that optogenetic activation of dopamine D3 receptor expressing neurons in the ventral striatal islands of Calleja robustly induces orofacial grooming in mice. Using this optogenetic manipulation, here we demonstrate that observer mice exhibit social preference for mice that groom more regardless of biological sex. Moreover, grooming-induced social attraction depends on volatile chemosensory cues broadcasted from grooming mice. Collectively, our study establishes self-grooming as a means of promoting social attraction among mice via volatile cues, suggesting an additional benefit for animals to allocate a significant amount of time to this behavior.

## Introduction

Self-directed grooming is essential for hygiene maintenance, thermoregulation, de-arousal, and stress reduction, and not surprisingly, animals allocate significant time to this behavior (1, 2). Although self-grooming is often conceptualized as a solitary or asocial behavior, it is implicated in social communication among conspecifics (3–7). For instance, ground squirrels engage in self-grooming to deescalate agonistic encounters during territorial disputes (7), and female meadow voles appear to be more attracted to males who groom (3, 5). These lines of evidence suggest that grooming broadcasts sensory cues that can influence behaviors among other nearby animals. However, direct evidence supporting this notion is still lacking, partly due to unpredictability of spontaneous grooming in experimental animals.

Recent advances in the understanding of neurobiological control of self-grooming make it possible to induce this behavior in laboratory mice via optogenetic manipulations of neuronal activity of specific cell types (8–10). Notably, the islands of Calleja (IC), clusters of densely-packed granule cells situated mostly in the olfactory tubercle (OT; also called tubular striatum (11)), contribute to a ventral striatal circuit that is involved in grooming control. The IC granule cells are characterized by expression of the dopamine D3 receptor, and optogenetic activation of these neurons reliably induces orofacial grooming (i.e., Phase I to III nose-face-head grooming without Phase IV body licking (12, 13)) in a temporally controllable manner (10).

Using this optogenetic manipulation, we can induce orofacial grooming in mice with temporal precision and directly address previously unanswered questions: 1) does orofacial grooming help mice attract conspecifics, 2) whether such attraction shows sexual dimorphism, and 3) what sensory channel conveys the grooming signal from sender to receiver. Our results demonstrate that observer mice spend more time investigating mice that groom more regardless of biological sex, and that such attraction is mediated by orofacial secretions from grooming mice and requires functional main olfactory epithelia of recipient mice. Overall, this study establishes self-grooming as a means of promoting social attraction via chemosensory communication.

## Results

### Mice showing more grooming attract nearby individuals regardless of biological sex

To investigate the potential role of self-grooming in social behaviors, we took advantage of an experimental approach which induces grooming with both reliability and temporal precision (10). The IC in the ventral striatum (mainly in the OT) are clusters of densely-packed, granule cells expressing the dopamine D3 receptor (**Fig. 1A, B, and Supplemental Video 1**). Optogenetic activation of these D3 neurons via an optical fiber implanted in the OT of double transgenic D3-Cre/ChR2-EYFP (or D3-ChR2 in brief) mice robustly induces orofacial grooming (**Fig. 1B, C, and Supplemental Video 2; (10)**). Since most OT/IC D3-ChR2 neurons fired at a maximum rate of 20 Hz upon current injection and faithfully followed 20 Hz blue light stimulation in brain slices (**Fig. 1B; (10)**), we used 20 Hz stimulation for all behavioral experiments. A three-chamber apparatus was used to assess social preference (**Fig. 1D)**. Two D3-ChR2 mice (same-sex littermates) with an optical fiber implanted in the OT were placed in each of the two side chambers under a cup (M1 and M2 in **Fig. 1D**). These two mice served as counterparts that groomed more or less, stimulated by blue or green light, respectively, while each observer mouse (M_ob_; unoperated) was placed in the center of the middle chamber and its behavior was recorded for 10 min (**Fig. 1D**). To prevent exhaustion of firing or neurotransmitter release of D3-ChR2 neurons, the light alternated between ON (10 s) and OFF (50 s) during the 10 min test. While the grooming time during green light stimulation was not significantly different from spontaneous grooming when light was off, blue light reliably elicited more grooming (7-8 sec/10 sec stimulation) than green light during the entire session **(Fig. 1E** and **Supplemental Video 3)**. During a 10 min test period for each observer mouse, blue light-stimulated mice exhibited a total of 120-140 sec orofacial grooming (light-induced plus spontaneous), which more than doubled the grooming time in green light-stimulated counterparts (**Fig. 1F**).

**Figure 1.**
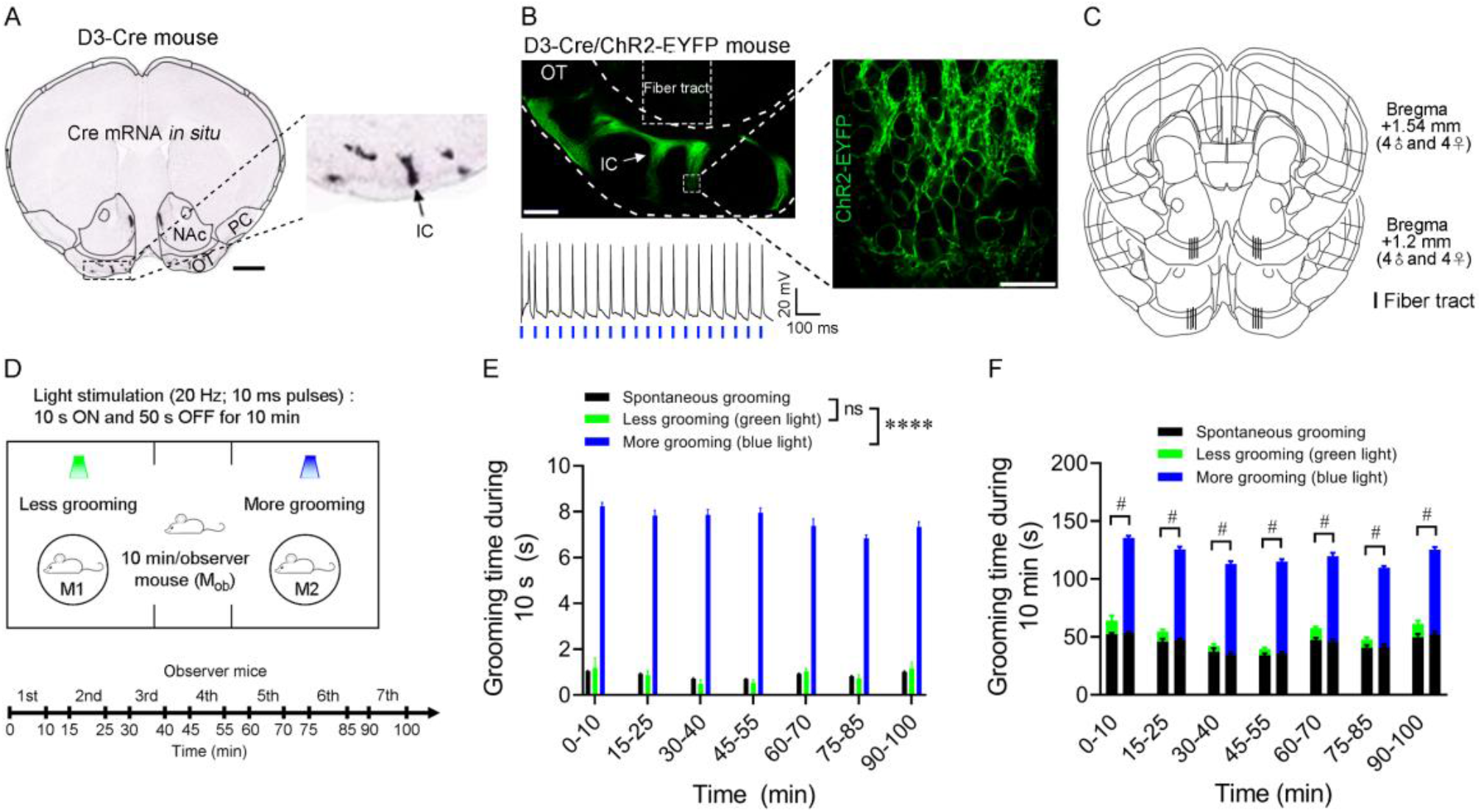
Paradigm for monitoring social preference during reliable induction of self-grooming. A. A coronal section across the ventral striatum showing *in situ* hybridization of Cre mRNA in a D3-Cre mouse. Image credit: Allen Institute (Allen Mouse Brain Connectivity Atlas: http://connectivity.brain-map.org/transgenic/experiment/304166273). Scale bar = 1 mm. Inset: an enlarged view of dotted rectangle area in left panel. Arrow denotes the IC. PC, piriform cortex. NAc, nucleus accumbens. OT, olfactory tubercle. IC, islands of Calleja. B. D3-ChR2 neurons are densely packed in the IC. Left, a representative image (coronal section) showing the IC and the optical fiber tract (upper), as well as firing of an IC D3-ChR2 neuron upon laser stimulation at 20 Hz (lower). Scale bar = 200 µm. Right, an enlarged image of the IC (dotted rectangle area in left panel). Scale bar = 20 µm. C. Coronal brain panels showing optical fiber placements in D3-ChR2 mice. D. Upper, schematic showing the behavioral strategy in the three-chamber apparatus. Blue light stimulation of OT D3-ChR2 neurons induced robust self-grooming (More grooming) while green light with the same stimulation parameters produced less (Less grooming). Lower, schematic depicting experimental timeline of the test. Ten min test for each observer mouse (M_ob_) with a 5 min interval between two consecutive mice. E-F. Comparison of total grooming time in 10 s (E) and 10 min (F) between spontaneous and light-stimulation conditions in an entire session. Data are shown as the mean ± s.e.m. n = 16 mice. **** or #, p < 0.0001, and ns, not significant. Results of statistical analyses are included in Supplemental Table 2.

We performed the initial social preference tests with two female mice stimulated by blue (more grooming) or green light (less grooming) under customized cups in the two side chambers, allowing emission of several possibly salient sensory cues (visual, auditory, and olfactory) **(Fig. 2A1)**. Each observer mouse (male) was placed in the middle chamber and the time it spent in each of the side chambers was recorded during 10 min, which was used to calculate the social preference index (see Methods for details; the total duration of stay for each observer mouse for all figures included in Supplemental Table S1). An index value of 0 indicates no preference and a positive or negative value indicates preference of an observer mouse toward one mouse or the other (**Fig. 2A2**). A control session (no light stimulation) with the same mice was performed 24 h prior to the light stimulation session. Compared to the no light control, in which the observer mice did not show a preference, they exhibited social preference toward the mouse that received blue light stimulation (more grooming) (**Fig. 2A2**). Such preference was also reflected in an increase in the total investigation time and number of investigation bouts (**Supplemental Fig. S1A**). To test whether blue light-stimulated mice (more grooming) attract conspecifics via visual communication, we covered the cups by paper towels with numerous tiny holes (**Supplemental Video 3**). Observer mice showed similar preference toward mice showing more grooming even in the absence of visual cues from light-stimulated mice (**Fig. 2B1, 2B2, and Supplemental Fig. S1B**), suggesting that visual cues are dispensable for grooming-induced social attraction.

**Figure 2.**
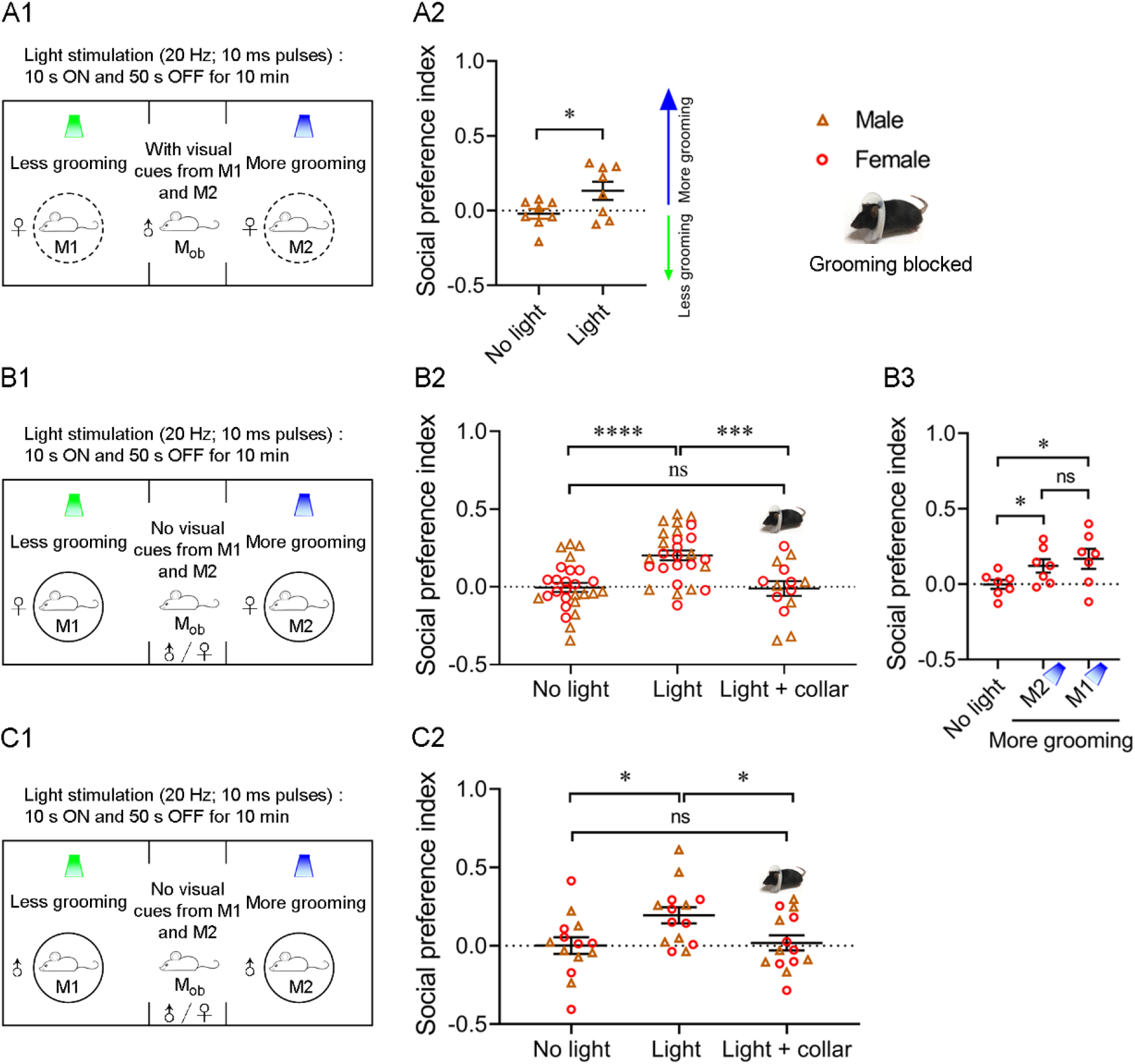
Observer mice show social preference for mice that groom more regardless of biological sex. A. Observer mice (n = 8) spent more time in the side of the mouse that groomed more (with visual cues). A1, schematic showing the behavioral strategy. A2, social preference index under no light and light stimulation conditions. B. Both male and female observer mice were attracted to the female mouse that groomed more (without visual cues). B1, schematic showing the behavioral strategy. B2, social preference index of observer mice under no light and light stimulation (with or without collar) conditions. B3, observer mice were always attracted to mice that groomed more. Light-stimulated mice were swapped by switching the blue/green light stimulation. C. Both male and female observer mice were attracted to male mice that groomed more. C1, schematic showing the behavioral strategy. C2, social preference index of observer mice under no light and light stimulation (with or without collar) conditions. For B2, n = 27 observer mice (13 males and 14 females). The observer mice in light + collar group were a subcohort of the mice tested. For B3, n = 7 observer mice (a subcohort of B2). For C2, n = 14 observer mice (7 males and 7 females). Data are shown as the mean ± s.e.m. * p < 0.05, ** p < 0.01, **** p < 0.0001, and ns, not significant. Raw data for calculating the social preference index are included in Supplemental Table S1 and results of statistical analyses in Supplemental Table S2.

We also examined whether grooming mice make audible calls and/or ultrasonic vocalizations by recording sounds up to 96 kHz. Grooming mice did not produce robust vocalizations, even though we were able to record spontaneous audible calls (**Supplemental Fig. S2A and Audio 1**). For some grooming strokes, we did record associated sounds, which appeared to result from physical contacts of the optical fiber tether and the recording chamber, rather than grooming *per se* (**Supplemental Fig. S2B and Audio 2**), as no reliable sounds were associated with spontaneous orofacial grooming in mice without a tether (not shown). We did not further investigate whether these sounds could attract other mice, given the sufficiency of chemosensory communication in grooming-induced social attraction as we uncovered by the following experiments.

Across the animal kingdom, many displays of communication are presented uniquely by each sex. Therefore, we investigated whether grooming-induced social attraction is sexually dimorphic. For either male or female light-stimulated mouse pairs, the observer mice (both male and female) exhibited a significantly higher social preference index toward blue light-stimulated mice (more grooming) compared to no light control (**Fig. 2B1, 2B2, 2C1, 2C2, and Supplemental Fig. S1B, C**). These data indicate that mice displaying more orofacial grooming attract other mice regardless of their biological sex.

Control experiments were conducted to ensure that the observed social preference was indeed attributed to orofacial grooming. First, as described above, in each set of experiments, the same observer mice did not show preference when no light was delivered to the two mice in the side chambers. Second, when the two light-stimulated mice wore a collar, which prevented orofacial grooming, social preference toward blue light-stimulated mice (more grooming) was eliminated (**Fig. 2B2, C2**). Upon blue light stimulation, collar-wearing mice still attempted grooming motions, but their forepaws failed to reach the orofacial parts (**Supplemental Video 4**). Note that during the experiments, the observer mice could not see light-stimulated mice under the covered cups. This experiment also ruled out the possibility that optogenetic activation of OT D3 neurons directly increases the secretion of volatile chemosensory cues (see below). Third, when the mice receiving blue or green light were swapped, or when the chamber assignment was switched, the observer mice always showed preference toward blue light-stimulated mice (more grooming) (**Fig. 2B3**). Fourth, the time that the observer mice spent in the side of blue light-stimulated mice was independent of the testing order (**Supplemental Fig. S3**), suggesting that orofacial grooming of blue light-stimulated mice was effective in attracting conspecifics throughout the entire session. Furthermore, we quantified the self-grooming behavior conducted by the observer mice when they were near light-stimulated mice and did not find a significant difference between the two sides (**Supplemental Fig. S1D)**, suggesting that grooming in one mouse does not promote self-grooming in nearby mice.

### Grooming-induced social attraction depends on chemosensory communication

Since olfactory cues are pivotal for social communication in mice, we sought to test the role of the olfactory system in orofacial grooming-induced social attraction by rendering the observer mice hyposmic. We intraperitoneally injected saline (as control) or methimazole in the observer mice and performed behavioral tests four days later (**Fig. 3A**). Methimazole treatment ablated the main olfactory epithelium (MOE) but left the vomeronasal epithelium intact **(Supplemental Fig. S4)**. Consistent with reduced perception of volatile odors (14), methimazole-treated observer mice decreased both the total time and number of bouts investigating light-stimulated mice compared to the saline controls (**Fig. 3C-E**). Nevertheless, methimazole-injected observer mice still moved around in the three-chamber arena, but they did not show social preference toward mice that groomed more as the saline-injected controls (**Fig. 3B**). This result suggests an essential role played by the MOE of the observer mice in this process, but due to potential off-target effects of methimazole and reduced social investigation, it is unclear whether there is a direct causal link between ablation of the MOE and lack of social preference.

**Figure 3.**
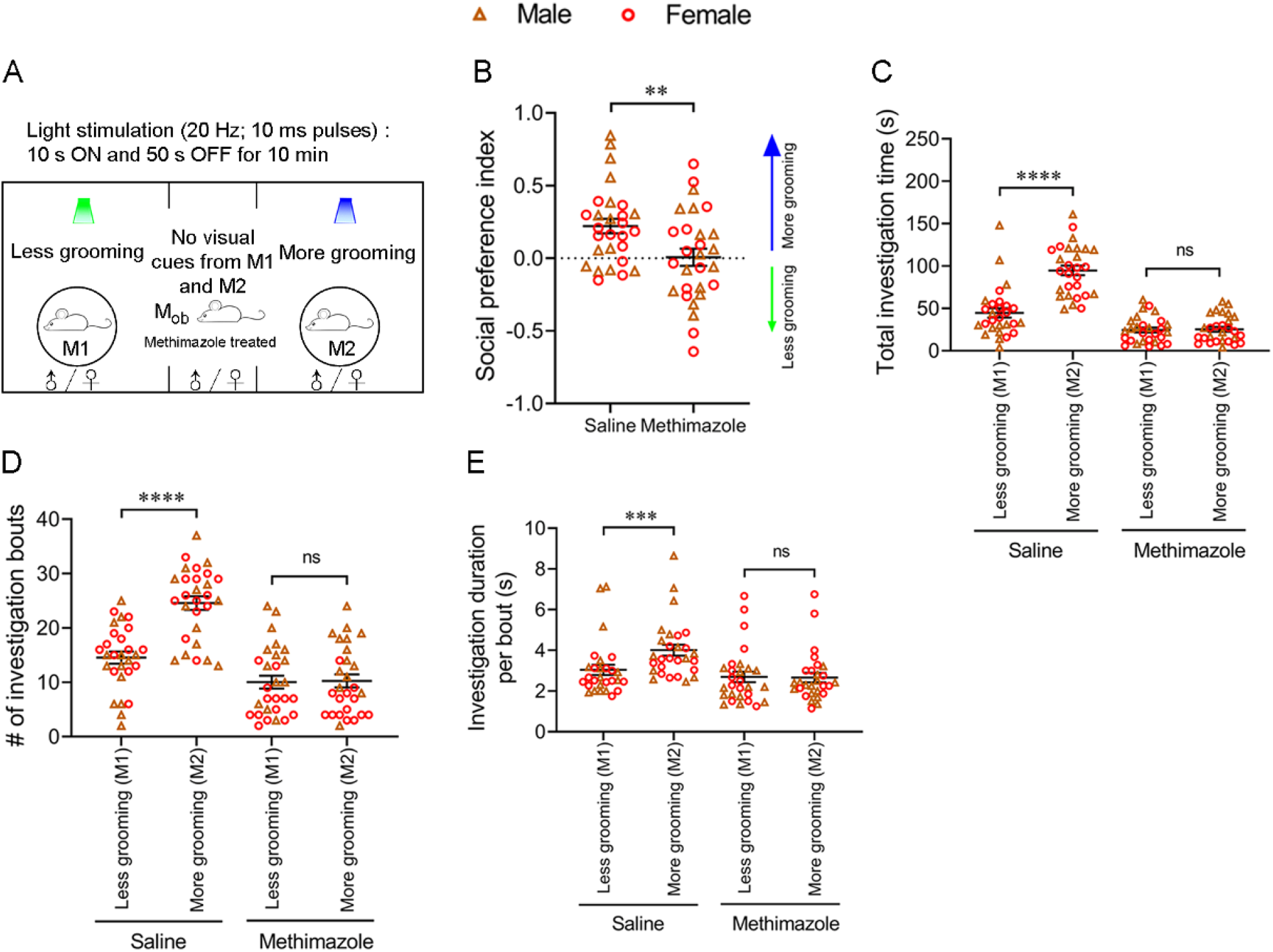
Social preference toward mice showing more grooming depends on functional main olfactory system of observer mice. A-B. Ablation of main olfactory epithelia of observer mice via methimazole treatment abolished their social preference toward blue light-stimulated mice (more grooming). A, schematic showing the behavioral strategy. B, social preference index of observer mice treated with saline (as control) or methimazole (hyposmic group). C-E. Methimazole treatment in observer mice reduced their investigation behavior. Quantification of total investigation time (C), number of investigation bouts (D), and investigation duration per bout (E), under the saline and methimazole treatment condition. n = 28 observer mice (14 males and 14 females) in each group. Data are shown as the mean ± s.e.m. ** p < 0.01, *** p < 0.001, **** p < 0.0001, and ns, not significant. Raw data for calculating the social preference index are included in Supplemental Table S1 and results of statistical analyses in Supplemental Table S2.

To provide direct evidence that self-grooming induces social attraction via orofacial chemical cues, we used mineral oil-moistened cotton swabs to sample the orofacial region (mouth, nose, cheek, and area surrounding eyes) of female mice that groomed more or less immediately after blue or green light stimulation **(Fig. 4A)**. The cotton swabs (each in a petri dish) were then placed under the cups to replace the light-stimulated mice in the previous experiments. Both male and female observer mice showed preference for the cotton swab from blue light-stimulated mice (more grooming) over the green light-stimulated mice (less grooming), in contrast to the control condition in which observer mice did not show preference for orofacial secretions of the same mice before they received light stimulation **(Fig. 4B)**. Similar findings were observed when the cotton swab experiments were conducted using orofacial secretions from male light-stimulated mice **(Fig. 4C, 4D)**. These results suggest that orofacial secretions from mice are broadcasted during grooming and are sufficient to attract conspecifics.

**Figure 4.**
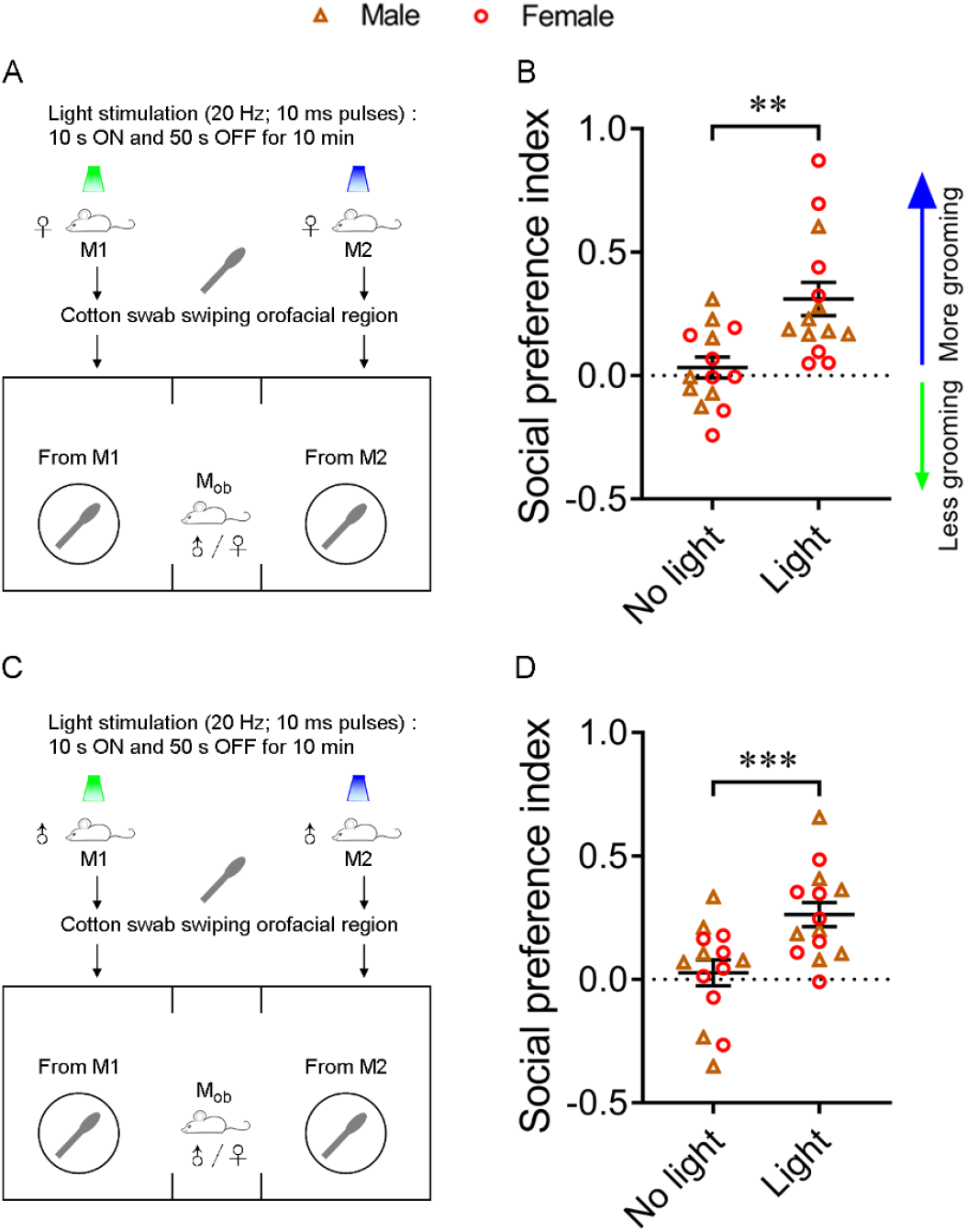
Observer mice spend more time investigating orofacial secretions from blue light-stimulated mice. A-B. Orofacial secretions from female blue light-stimulated mice attracted both male and female conspecifics. A, schematic showing the behavioral strategy. B, social preference index of observer mice under no light and light condition. C-D. Orofacial secretions from male blue light-stimulated mice attracted both female and male conspecifics. C, schematic showing the behavioral strategy. D, social preference index of observer mice under no light and light condition. n = 14 observer mice (7 males and 7 females). In B and D, orofacial secretions collected under the light condition were from the same mice under the no light condition. Data are shown as the mean ± s.e.m. ** p < 0.01 and *** p < 0.001. Raw data for calculating the social preference index are included in Supplemental Table S1 and results of statistical analyses in Supplemental Table S2.

## Discussion

In the present study, using an optogenetic approach to induce orofacial grooming in a reliable and controllable manner, we demonstrate in mice that self-grooming promotes social attraction and that, regardless of biological sex, mice are attracted to mice that groom more. This effect is predominantly mediated by orofacial secretions emitted as volatiles during self-grooming and perceived via the recipient’s main olfactory system. This work complements observational studies among other types of animals in the field and extends those observations by providing insights into a causal sensory channel involved in intra-specific communication via self-grooming.

Self-grooming allows an animal to emit rich sensory cues which potentially affect the behaviors of nearby recipients. Grooming without doubt is visually observable and thus a likely channel whereby it is perceived by an observer would be visual. In contrast, in the present experiments, we uncovered in mice that visual and even auditory cues may be dispensable for grooming-induced social attraction because the effect persists without visual cues (**Fig. 2**) and robust audible calls are not emitted during grooming (**Supplemental Fig. S2**). Compared to visual and auditory signals, chemosensory cues offer the unique ethological advantage of communication over time and space. In fact, rodents heavily rely on chemosensory cues for their social behaviors (e.g., (15–19)). Through self-grooming, individuals can further the volatility of their bodily scents, potentially attracting conspecifics (3–6). Rodents use multiple chemosensory organs including the MOE and vomeronasal organ to detect social cues (15–19). While the vomeronasal organ is involved in communication via pheromones which are largely nonvolatile, the MOE receives and supports the processing of mainly volatile odors (20, 21). Consistent with this notion, methimazole treatment, which ablated the main olfactory epithelium but not the vomeronasal organ in the recipients (**Supplemental Fig. S4**), abolished their social preference for mice showing more grooming (**Fig. 3**). Although methimazole reduced social investigation in general and might have off-target effects, our findings support that the main olfactory system of the recipient is required for social preference toward mice showing more grooming. We can’t not completely rule out the possibility that spontaneous grooming may not increase the secretion of volatile chemosensory cues. But we think this scenario is unlikely, because the orofacial grooming induced by optogenetic activation of OT D3 neurons is indistinguishable from spontaneous grooming and the increased grooming time (approximately from 10% to 20% of the total time within each 10 min) is still within the physiological range.

This finding is not without precedent. It has been proposed that grooming animals may emit volatile cues to communicate their sex, identity, and reproductive status and thereby attract nearby opposite-sex conspecifics (3–6). However, to what extent self-grooming serves as a sexually dimorphic communication signal remains to be addressed. Male voles spend more time self-grooming when they are exposed to odors of females, and are more responsive to grooming females (3, 5, 6). However, the frequency and duration of self-grooming of male voles do not predict mating success (22). Here we demonstrate that mice that groom more attract conspecifics regardless of sex (**Fig. 2**), suggesting that, in mice, self-grooming provides broadly-appealing social communicative signals. The discrepancies between these studies may mainly result from two reasons. First, secretions released from orofacial glands (e.g., salivary and Harderian glands) and their effects on the recipients may vary in different species. Even in the same species, the secretions may vary under different contexts, hormonal statuses, and experimental conditions. For example, in Mongolian gerbils, Harderian secretions released during self-grooming are enhanced when the animals are exposed to cold temperatures (23). Second, in rodents, a complete grooming bout consists of a syntactic chain that progresses sequentially from nose-face-head (phase I-III) grooming to body licking (phase IV) (12, 13). Bodily secretions induced by distinct grooming phases probably vary, which may lead to different behavioral effects in the recipients even in the same species. The optogenetic approach used in our study induces orofacial grooming but not body licking (10). Body licking involves licking of the genital area, which is correlated with sexual behavior in rats (24), probably due to grooming-induced release of sex-specific materials from genital glands.

We found that cotton swabs rubbed against the orofacial parts from mice showing more grooming are sufficient to attract the recipient mice (**Fig. 4**), suggesting that self-grooming may lead to release of compounds from specifically the orofacial glands. Consistent with this finding, grooming-induced social attraction was absent when direct contacts of the forepaws with orofacial parts were prevented by a collar (**Fig. 2**). Orofacial grooming releases and spreads compounds from various glands such as salivary and Harderian (25–27). These compounds can serve as social communicative signals in rodents, including mouse (28), Mongolian gerbil (29–31), and golden hamster (32). Our study reveals that orofacial grooming attracts other mice regardless of sex, but it does not rule out the possibility that orofacial grooming also releases sex-specific signals. In fact, the Harderian gland in golden hamster exhibits pronounced sexual dimorphism in histology and products, and males are more attracted to fresh smears of Harderian gland from females than males (33). In addition, Harderian gland secretions from male Mongolian gerbils contribute to the proceptive behaviors of estrous females (29). Future studies are warranted to identify the specific types or combinations of orofacial secretions involved in grooming-induced general social attraction versus sexual attraction.

## Methods

### Animals

The transgenic D3-Cre line (STOCK Tg(Drd3-cre)KI198Gsat/Mmucd, RRID:MMRRC_031741-UCD) was obtained from the Mutant Mouse Resource and Research Center (MMRRC) at University of California at Davis, an NIH-funded strain repository, and was donated to the MMRRC by Nathaniel Heintz, Ph.D., The Rockefeller University, GENSAT and Charles Gerfen, Ph.D., National Institutes of Health, National Institute of Mental Health. The D3-Cre line was crossed with the Cre-dependent channelrhodopsin 2 (ChR2) line (JAX Stock No: 024109 or Ai32 line: Rosa26-CAG-LSL-ChR2(H134R)-EYFP-WPRE) (34) to generate D3-Cre/ChR2-EYFP (or D3-ChR2) mice. Mice were maintained in temperature- and humidity-controlled animal facilities under a 12 h light/dark cycle with food and water available *ad libitum*. Both male and female mice (2-3 months old) were used. Mice were group-housed until the surgery of intra-cranial implantation and singly-housed afterwards. All experimental procedures were performed in accordance with the guidelines of the National Institutes of Health and approved by the Institutional Animal Care and Use Committee of the University of Pennsylvania.

### Stereotaxic surgery and optical fiber implantation

Mice were anesthetized with isoflurane (~3% in oxygen) and secured in a stereotaxic system (Model 940, David Kopf Instruments). Isoflurane levels were maintained at 1.5-2% during the surgery. Body temperature was monitored and maintained at 37 °C with a heating pad connected to a temperature control system (TC-1000, CWE Inc.). Local anesthetic (bupivacaine, 2 mg/kg, s.c.) was applied before skin incision and hole drilling on the dorsal skull. In order to target the IC in the OT, which is a large structure, two sets of coordinates from bregma were used: anteroposterior (AP) 1.2 (or 1.54) mm; mediolateral (ML) ±1.1 (or 1.15) mm; dorsoventral (DV), −5.5 (or −5.0) mm. The results were combined since we did not observe significant differences. A cannula (CFMC14L10-Fiber Optic Cannula, Ø2.5 mm Ceramic Ferrule, Ø400 µm Core, 0.39 NA; Thorlabs, Newton, NJ), customized to 6 mm length, was placed in the OT at the same coordinates as described above and fixed on the skull with dental cement. D3-ChR2 mice were returned to home cage for recovery for one week before behavioral tests. Only mice showed robust grooming behavior upon blue light stimulation were used in behavioral tests and optical fiber placements near the IC were verified *post-mortem* for all operated mice.

### *In vivo* optical stimulation and behavioral assays

Behavioral tests were performed during the light cycle (9:00 am - 12:00 pm). Mice were acclimated to the testing room at least 1 h before the tests. In experiments using collars, mice were habituated with wearing the collar 2 h per day for 3 consecutive days. Before each test, mice were briefly anesthetized via isoflurane and a flexible optic tether was coupled to the implanted fiber stud with a mating sleeve (Thorlabs Inc.).

A pair of D3-ChR2 mice (same-sex littermates) with optical fiber implanted in the OT were placed in each of the two side chambers in a three-chamber apparatus (30 cm × 15 cm × 20 cm) and covered under plastic cups. The wall of the cups had parallel vertical cuts (~10 mm in width with ~15 mm between two cuts) so that the observer mice could visualize the mice in the side chambers. In experiments without visual cues, the cups were wrapped in paper towels punched with numerous tiny holes (~1 mm in diameter with ~5 mm between two holes) for ventilation. Each observer mouse (typically 7 in an entire session) was placed in the center of the middle chamber of the three-chamber apparatus at the beginning of each test. The locations of blue and green light-stimulated mice were counterbalanced across different tests to avoid any side bias. Each observer mouse was subjected to two tests with an interval of 24 h, in which the pair of D3-ChR2 mice received no light (control condition) or blue/green light stimulation (experimental condition). For light stimulations, one mouse was stimulated with blue laser (473 nm) and the other one with green laser (532 nm) using the same parameters (10-15 mW; 20 Hz with 10 ms pulses). During the 10 min test for each observer mouse, the light stimulation was delivered in a protocol of 10 s ON and 50 s OFF. In experiments where self-grooming was blocked, light-stimulated mice wore a collar to prevent their forepaws from touching orofacial parts. In a subset of experiments, 24 h after the initial test, the location as well as the light pairing were switched for the two mice in the side chambers to exclude potential biases of the side and mouse. For methimazole treatment, observer mice were intraperitoneally injected with either saline (as control) or methimazole (75 mg/kg), and behavioral tests were conducted four days post injection. The behavioral tests were video-taped via a webcam (30 frames/sec) and analyzed post hoc using the Any-Maze software by experimenters who were blinded to the experimental conditions. According to a standardized protocol (35), the social preference index was calculated as (T2-T1)/(T2+T1), where T2 and T1 were the time of an observer mouse spent (i.e., total duration of stay) in each side chamber with the blue (more grooming) or green (less grooming) light-stimulated mouse, respectively. An investigation bout was defined from the time when an observer mouse started to sniff the cup covering a light-stimulated mouse (the nose was within ~0.5 cm from the cup) to the time when the observer mouse turned away. The total investigation time was the sum of the time the observer mouse spending in all investigation bouts toward one cup during a 10 min test.

### Collection of orofacial secretions from light-stimulated mice

Mice were first subjected to either blue or green light stimulations as aforementioned for 10 min. Secretions were immediately collected from the orofacial region (including the mouth, nose, cheek, and area surrounding the eye) using Q-tip cotton swabs that were moistened by mineral oil. For each mouse, a cotton swab was swiped against the orofacial region a total of 15-20 times. The cotton swabs were then placed in petri dishes (diameter: 6 cm), covered by the two cups in the two side compartments of the three-chamber apparatus. Observer mice were then placed at the center of the middle chamber at the beginning of each test, and their activities were videotaped for 10 min for post hoc calculation of the social preference index. The placement of the petri dish was counterbalanced between different tests to avoid potential side bias.

### Audio recording

Audio recording of blue laser induced self-grooming was performed in a sound-attenuated chamber. Within this chamber, a condenser ultrasound microphone (CM16/CMPA, Avisoft Bioacoustics) was fixed above a clean cage. Mice were habituated to the sound attenuated chamber and underwent 4 laser stimulation trials using parameters described above (20 Hz 10 ms pulses for 10 s) with a 5 min interval between stimulations. Acoustic data were acquired at 192 kHz to capture potential audible and ultrasonic sounds related to self-grooming. Data were acquired, visualized, and quantified using Raven Pro v1.4 software (the Cornell Lab of Ornithology).

### *Ex vivo* electrophysiological recordings

Whole-cell patch-clamp recordings were performed as we described previously (10). Mice were deeply anesthetized with ketamine-xylazine (200 and 20 mg/kg body weight, respectively) and decapitated. The brain was dissected out and immediately placed in ice-cold cutting solution containing (in mM) 92 N-Methyl D-glucamine, 2.5 KCl, 1.2 NaH_2_PO_4_, 30 NaHCO_3_, 20 HEPES, 25 glucose, 5 Sodium L-ascorbate, 2 Thiourea, 3 Sodium Pyruvate, 10 MgSO_4_, and 0.5 CaCl_2_; osmolality ~300 mOsm and pH ~7.3, bubbled with 95% O_2_-5% CO_2_. Coronal sections (250 µm thick) containing the OT were cut using a Leica VT 1200S vibratome. Brain slices were incubated in oxygenated artificial cerebrospinal fluid (ACSF in mM: 124 NaCl, 3 KCl, 1.3 MgSO_4_, 2 CaCl_2_, 26 NaHCO_3_, 1.25 NaH2PO_4_, 5.5 glucose, and 4.47 sucrose; osmolality ~305 mOsm and pH ~7.3, bubbled with 95% O2-5% CO_2_) for ~30 min at 31°C and at least 30 minutes at room temperature before use. For recordings, slices were transferred to a recording chamber and continuously perfused with oxygenated ACSF. D3-ChR2 cells were visualized through a 40X water-immersion objective on an Olympus BX61WI upright microscope equipped with epifluorescence.

Whole-cell patch-clamp recordings were controlled by an EPC-10 amplifier combined with Pulse Software (HEKA Electronik) and analyzed using Igor Pro 6 (Wavematrics). Recording pipettes were made from borosilicate glass with a Flaming-Brown puller (P-97, Sutter Instruments; tip resistance 5-10 MΩ). The pipette solution contained (in mM) 120 K-gluconate, 10 NaCl, 1 CaCl_2_, 10 EGTA, 10 HEPES, 5 Mg-ATP, 0.5 Na-GTP, and 10 phosphocreatine. Light stimulation was delivered through the same objective via pulses of blue laser (473 nm, FTEC2473-V65YF0, Blue Sky Research, Milpitas, USA) at 20 Hz with 10 ms pulse.

### Immunostaining and confocal imaging

Mice were transcardially perfused with 4% paraformaldehyde (PFA) in fresh phosphate buffered saline (PBS). For *post-mortem* verification of optical fiber placement, brains were post fixed in 4% PFA overnight at 4 °C, then transferred into PBS. Coronal slices at 100 μm thick were prepared using a Leica VT 1200S vibratome. The slices were treated with glycerol in PBS (volume ratio 1:1) for 30 min followed by glycerol in PBS (volume ratio 7:3) for 30 min before being mounted onto superfrost slides (Fisher Scientific) for imaging. For immunostaining of the nasal tissues, the heads were post fixed 4% PFA overnight at 4 °C, and then decalcified in 0.5 M EDTA (pH 8.0, ethylenediaminetetraacetic acid) for four days and infiltrated in a series of sucrose solutions before being embedded in OCT. The frozen tissues were cut into 20 μm coronal sections on a cryostat. After antigen retrieval in a 95 °C water bath for 10 min, the tissue sections were blocked for 30 min in 0.3% Triton X-100 in PBS with 3% bovine serum albumin, and then incubated at 4 °C overnight in the same solution with the primary rabbit anti-OMP (olfactory marker protein; 1:500, O7889 from Sigma). Immunofluorescence was achieved by reaction with the secondary antibody donkey anti-rabbit-488 (A21206 from Molecular Probes, Invitrogen) at 1:200 for one hour. Tissues were washed in 0.3% Triton X-100 in PBS and mounted in Vectashield (Vector Laboratories). Fluorescent images were taken under a SP5/Leica confocal microscope with LAS AF Lite software.

### Statistical analyses

Graphs are created in GraphPad Prism and assembled in Adobe Photoshop. Normal distribution of datasets was tested via Shapiro-Wilk tests and parametric or non-parametric statistical tests were used accordingly.

## ACKNOWLEDGEMETNS

We thank Steven Simmons and Amelia Eisch for their help in setting up the sound recording system, and Usuy David Leon Tolosa for his assistance in analyzing some of the behavioral videos. This work was supported by the National Institutes of Health R01NS117061, R01DA049545, and R01DA049449 to M.M. and D.W.W., R01DC006213 to M.M., R21DC019193 to J.P.B., and F31MH124372 to E.J.

## Supplemental Figures

**Supplemental Figure S1.**
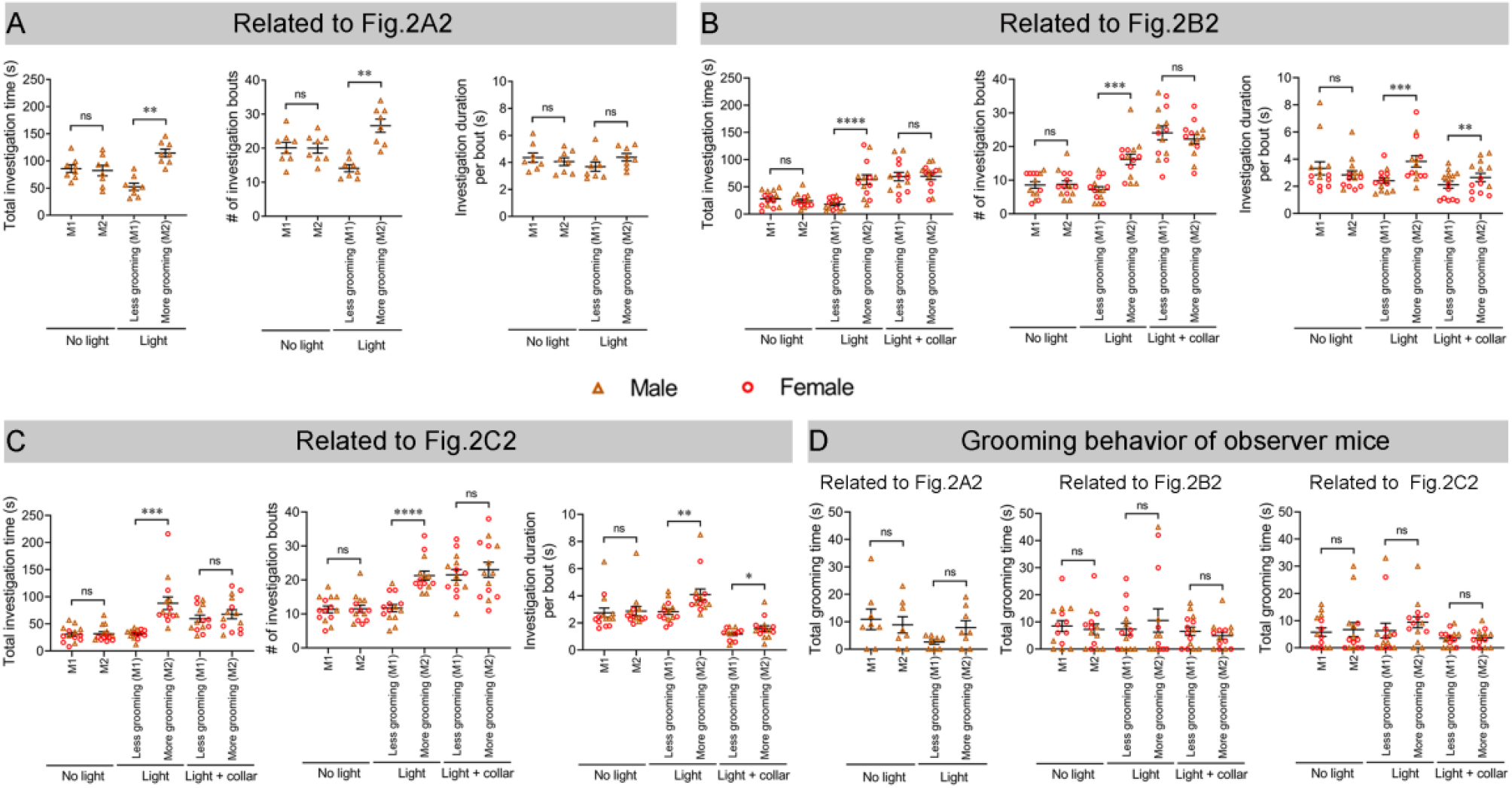
Observer mice spend more time investigating mice that groom more. A. Mice stimulated by blue (more grooming) or green light (less grooming) in uncovered cups (with visual cues). Investigation behavior of observer mice (n = 8 male) with female light-stimulated mice. B-D. Mice stimulated by blue (more grooming) or green light (less grooming) in covered cups (without visual cues). B, investigation behavior of observer mice (n = 13 males and 14 females) with female light-stimulated mice. C, investigation behavior of observer mice (n = 7 males and 7 females) to male light-stimulated mice. D, observer mice show similar durations of self-grooming near blue or green light-stimulated mice. Left, light-stimulated mice in uncovered cups (with visual cues). Self-grooming behavior of observer mice (n=8 male) near female blue or green light-stimulated mice. Middle and right, light-stimulated mice in covered cups (without visual cues). Middle, self-grooming behavior of observer mice (n = 13 males and 14 females) near female blue or green light-stimulated mice. Right, self-grooming behavior of observer mice (n = 7 males and 7 females) near male blue or green light-stimulated mice. Data are shown as the mean ± s.e.m. * p < 0.05, ** p < 0.01, *** p < 0.001, **** p < 0.0001, and ns, not significant. Results of statistical analyses are included in Supplemental Table S2.

**Supplemental Figure S2.**
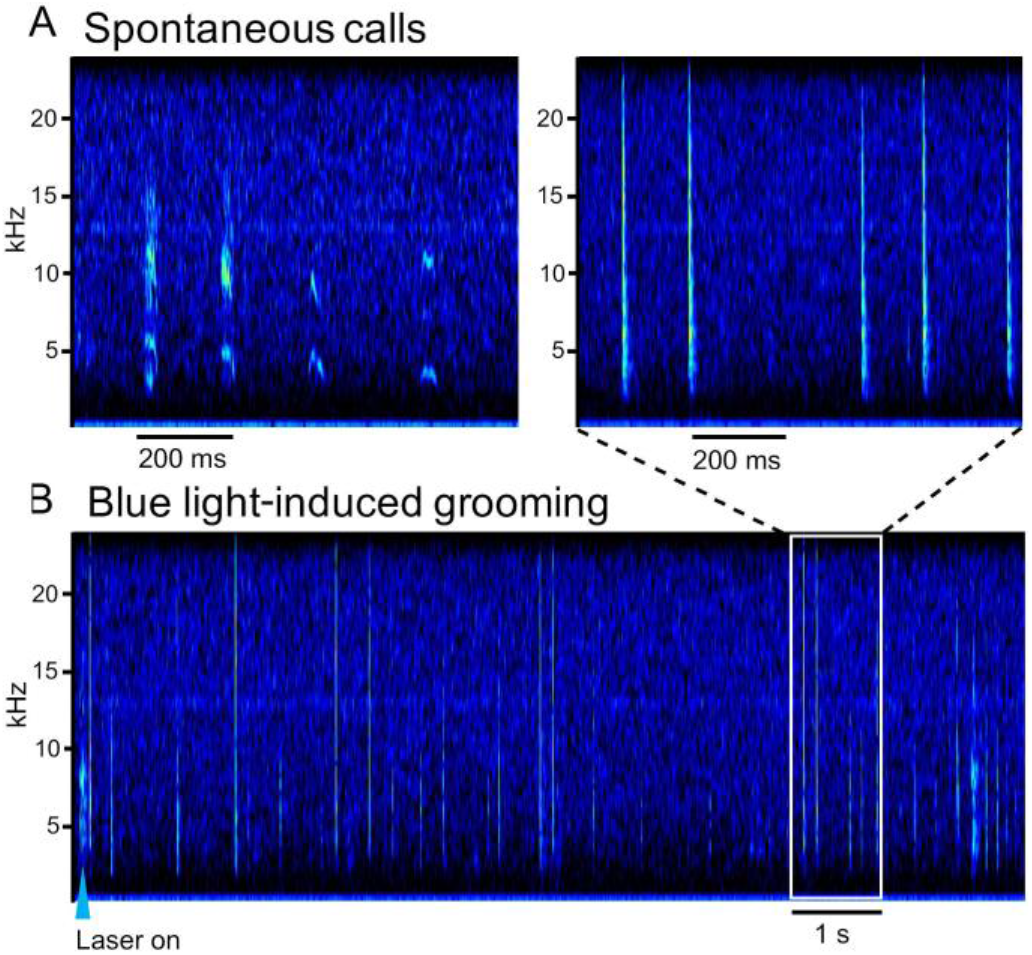
Mice do not emit discernable vocalizations during self-grooming. Sonograms of spontaneous audible calls (A) and sounds associated with grooming strokes (B). No obvious ultrasonic vocalizations between 20 to 96 kHz in these recordings (not shown). Inset, an enlarged view of the rectangle area in (B). Whereas spontaneous calls, as expected, were restricted within frequency bands, self-grooming was associated with broad-band acoustic events which corresponded in time to movement of the fiber optic tether. Similar results were obtained from 6 mice recorded.

**Supplemental Figure S3.**
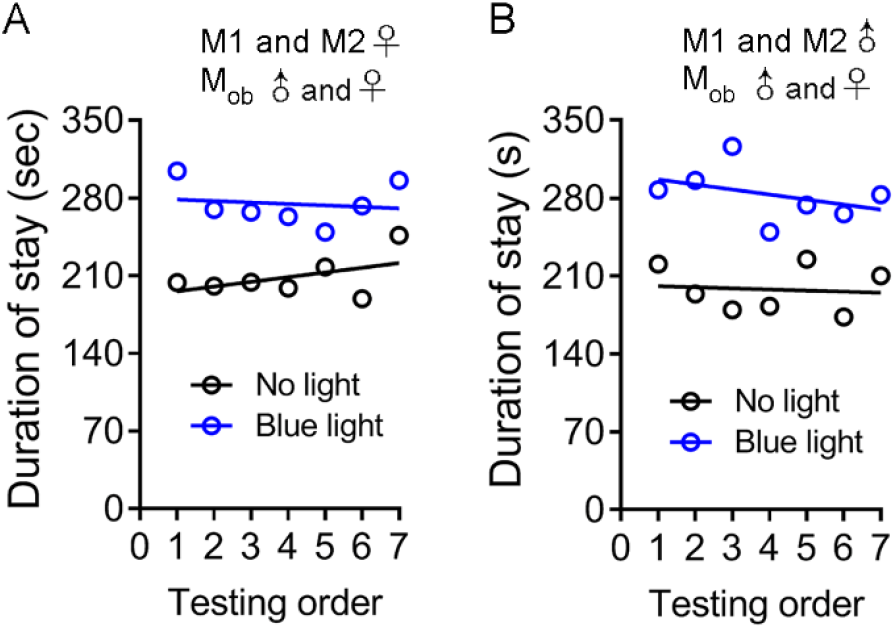
The testing order of observer mice does not impact the time spent in the side of blue light-stimulated mouse. The duration of stay in the side of blue light-stimulated mouse (more grooming) is plotted against the testing order of observer mice. A. female D3-ChR2 mice with no light or blue light stimulation in the OT. B. male D3-ChR2 mice with no light or blue light stimulation in the OT. Each data point is an average of four observer mice (two male and two female).

**Supplemental Figure S4.**
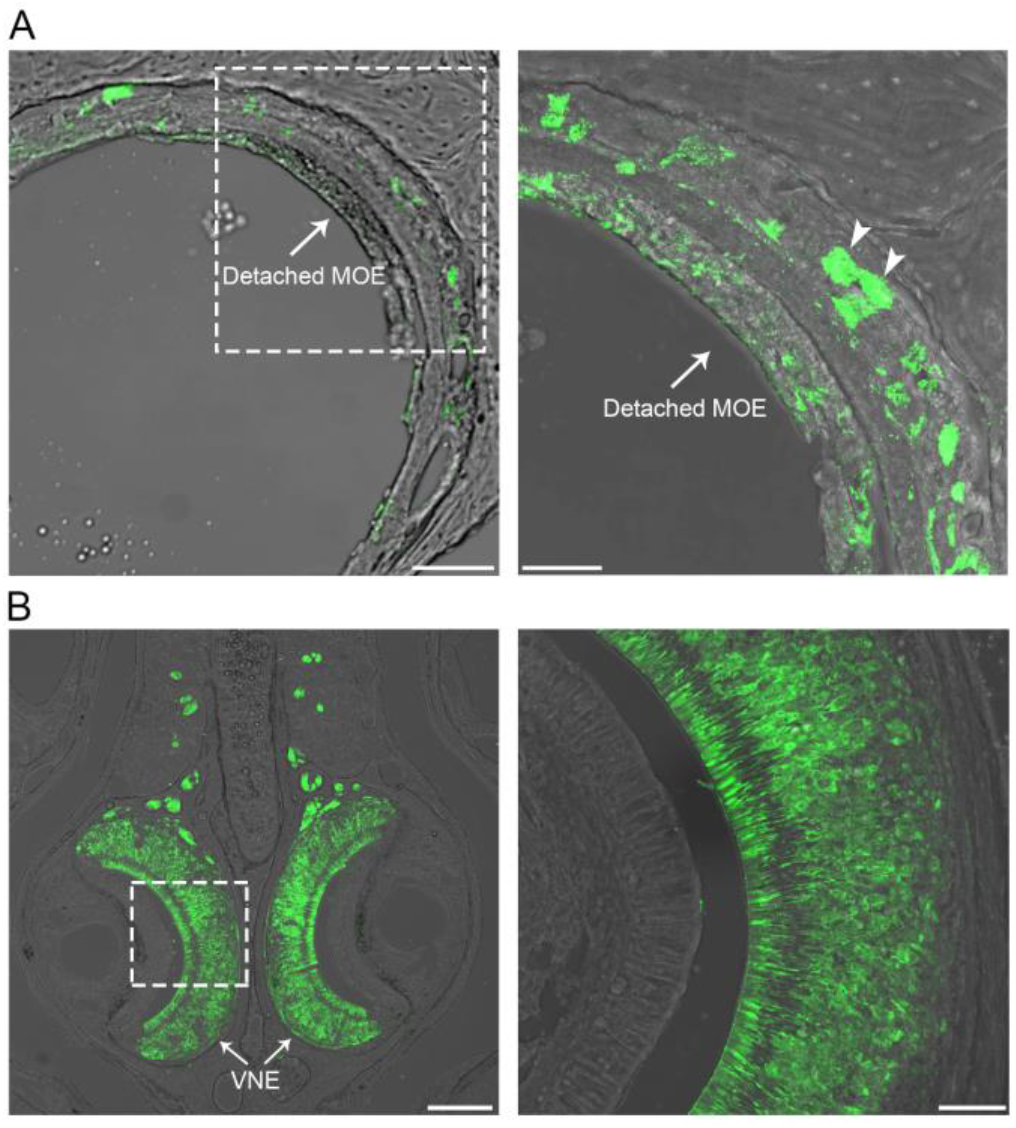
Methimazole treatment ablates the main olfactory epithelium but leaves the vomeronasal epithelium intact. Four days post methimazole treatment, coronal sections of the nose were processed for immunoreactivity to the olfactory marker protein (OMP), which labels mature olfactory sensory neurons. A. Left, a representative image showing the detached main olfactory epithelium (MOE) without obvious OMP+ olfactory sensory neurons (OSNs). Right, an enlarged view of the dotted rectangle area in the left panel. Arrowheads denote OSN axon bundles. Scale bars = 100 (left) and 50 µm (right). B. Left, a representative image (coronal section) showing the intact vomeronasal epithelium with abundant OMP+ sensory neurons. Right, an enlarged view of the dotted rectangle area in the left panel. Scale bars = 200 (left) and 50 µm (right). MOE, main olfactory epithelium; VNE, vomeronasal epithelium. Similar results were obtained from 3 mice.

## Supplemental Videos and Audios

**Video 1**. Z-stack confocal images showing densely packed D3-ChR2 neurons in an island of Calleja in the OT.

**Video 2**. Blue light (but not green light) stimulation of OT D3-ChR2 neurons induces orofacial grooming.

**Video 3**. An observer mouse spends more time investigating a blue light-stimulated mouse (more grooming) than a green light-stimulated counterpart (less grooming) in a three-chamber apparatus.

**Video 4**. Orofacial grooming induced by blue light stimulation of OT D3-ChR2 neurons is prevented by a collar.

**Audio 1**. Spontaneous audible calls recorded from a D3-ChR2 mouse.

**Audio 2**. Sounds recorded during blue light stimulation-induced grooming in a D3-ChR2 mouse. These sounds likely result from physical contacts of the optical fiber tether and the recording chamber, rather than grooming *per se*.

